# Simple adjustment of the sequence weight algorithm remarkably enhances PSI-BLAST performance

**DOI:** 10.1101/092742

**Authors:** Toshiyuki Oda, Kyungtaek Lim, Kentaro Tomii

**Affiliations:** Artificial Intelligence Research Center, National Institute of Advanced Industrial Science and Technology (AIST), 2-4-7 Aomi, Koto-ku, Tokyo 135-0064, Japan; Biotechnology Research Institute for Drug Discovery, National Institute of Advanced Industrial Science and Technology (AIST), 2-4-7 Aomi, Koto-ku, Tokyo 135-0064, Japan

## Abstract

PSI-BLAST, an extremely popular tool for sequence similarity search, features the utilization of Position Specific Scoring Matrix (PSSM) constructed from a multiple sequence alignment (MSA). PSSM allows the detection of more distant homologs than a general amino acid substitution matrix does. An accurate estimation of the weights of sequences in an MSA is crucially important for PSSM construction. PSI-BLAST divides a given MSA into multiple blocks, for which sequence weights are calculated. When the block width becomes very narrow, the sequence weight calculation can be difficult.

We demonstrate that PSI-BLAST indeed generates a significant fraction of blocks having widths less than 5, thereby degrading the PSI-BLAST performance. We revised the code of PSI-BLAST to prevent the blocks from being narrower than a given minimum block width (MBW). We designate the modified application of PSI-BLAST as PSI-BLASTexB. When MBW is 25, PSI-BLASTexB notably outperforms PSI-BLAST consistently for three independent benchmark sets. The performance boost is even more drastic when an MSA, instead of a sequence, was used as a query.

Our results demonstrate that the generation of narrow-width blocks during the sequence weight calculation is a critically important factor that restricts the PSI-BLAST search performance. By preventing narrow blocks, PSI-BLASTexB remarkably upgrades the PSI-BLAST performance.

## Introduction

Sequence similarity search is an initial choice for the functional inference of unknown biological sequences, for which BLAST (Altschul et al. 1990) is widely used. BLAST uses an amino acid substitution matrix such as BLOSUM62 (Henikoff & Henikoff 1992) to score distances between amino-acid pairs. BLAST, however, fails to detect distant homologs, partially because the matrices only provide a fixed score for a given amino-acid pair, although we know that the preference of amino acids varies among homologs and among positions within homologs.

The multiple sequence alignment (MSA) of closely related homologous sequences detected by BLAST is expected to contain such homolog-specific and position-specific information. An MSA can be transformed into a position-specific scoring matrix (PSSM), which is a more sophisticated model for sequence similarity search than the substitution matrix because scores for amino-acids are modeled for individual positions. Iterative search methods including PSI-BLAST (Altschul et al. 1997) construct a PSSM from an MSA obtained from the previous search. Then such methods use the PSSM for another similarity search. It has been demonstrated that much more distant homologs can be detected by iterating these steps. Because of its usefulness and availability, many modifications have been proposed since PSI-BLAST was first published, including introduction of composition-based statistics, optimizing cache utilization, and revising pseudo-count strategy (Altschul et al. 2009; Aspnas et al. 2010; Schaffer et al. 2001). Overcoming the problem of “homologous over extension” also improves the PSI-BLAST accuracy (Gonzalez & Pearson 2010; Li et al. 2012). In this study, we describe that PSI-BLAST can be improved further by slightly changing the sequence weighting method.

Because sequences in public databases are highly biased into organisms which are medically and commercially important, and because they are easy to culture, it is crucially important to adjust amino acid observations in the MSA of homologous sequences before PSSM calculation. Sequence weight is a straightforward means of attaining such adjustment, where a sequence with more closely related counterparts in the MSA should be assigned a smaller weight. PSI-BLAST calculates the position-specific sequence weight (PSSW) using a procedure derived from the formula proposed (Henikoff & Henikoff 1994) as:

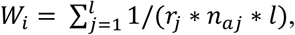

where *Wi* stands for the weight of *i*th sequence in a MSA, *r_j_* denotes the number of unique amino acids found at the position *j*, *l* signifies the length of the alignment, and *n_aj_* represents the number of amino acids *a* found at *j*. After sequences are weighted, the probability of *a* at *j* (*P_aj_*) is calculated as:

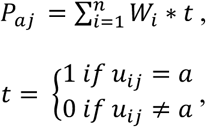

where *u_ij_* is the amino acid at *j* in the *i*th sequence.

This formula lacks the consideration of gaps. Simply put, gaps (including N-terminal and C-terminal gaps) can be treated as the 21st amino acid. An important problem of this approach is that the weights of gappy sequences in a gappy MSA will be underestimated. One can avoid this problem by considering an MSA subregion with few or no gaps for PSSW calculation. This is expected to be advantageous for dealing with MSAs constructed from local alignments that are likely to include many gaps. PSI-BLAST defines such blocks for individual positions. PSI-BLAST first selects a subset of sequences (a reduced MSA) in an MSA, such that no gap is included at a position of interest *j*. PSI-BLAST then collects starting and ending positions of all pairwise alignments between query and subjects in the reduced MSA to define the boundary of the block as the starting and ending positions closest to *j* (Altschul et al. 1997). This approach also has an important limitation: the block width might become extremely narrow, failing to reflect actual evolutionary information.

This study demonstrates that such narrow blocks are created during the PSSM construction of PSI-BLAST, which gives rise to inaccurate calculation of PSSW and PSSM, and which thereby drastically hampers the homology detection performance. We propose a simple method for better PSSW calculation, which boosts the performance of PSI-BLAST.

## Methods

### Narrow blocks result in wrong sequence weight calculation

Two artificial MSAs are presented in (Figure 1). The MSA in Figure 1A (MSA-A) is a subset of AAA ATPase MSA in the Pfam database (Finn et al. 2016). The MSA in Figure 1B (MSA-B) is identical to MSA-A except for 10th and 11th sequences, which were derived from the tenth sequence in MSA-A by dividing it into two pieces with an overlap at position 19. The two MSAs were converted to PSSMs (Supplemental Files 1 and 2, respectively) by PSI-BLAST search against a dummy database with “-in_msa”, “-num_iterations 1” and “-out_ascii_pssm” options.

**Figure 1.**
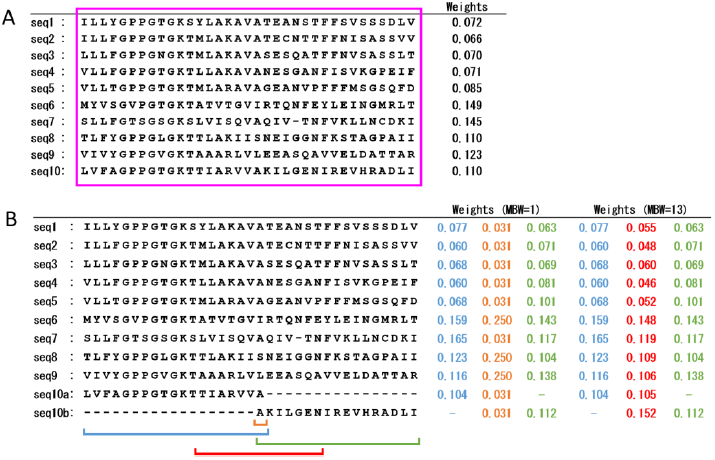
Examples showing the sequence weight calculation of PSI-BLAST and PSI-BLASTexB. (A) Sequence weights (shown on the right side) of all positions in the MSA were calculated from a single block covering the whole alignment. (B) PSI-BLAST divided the MSA into three blocks (blue, orange, and green) and calculated sequence weights for each block. Sequence weights calculated from the blocks are shown on the right side with the same color. For the orange block that is one aa long, PSI-BLASTexB extends the block such that the block width becomes MBW (red block). Weights calculated from the red block are also shown. See *Methods* for detailed procedures. “seq7” has no an amino acid at position 23. For that reason, the sequence weights of other sequences are calculated ignoring “seq7”.

We checked inner variables of PSI-BLAST to mark blocks on MSA-A and MSA-B (Fig. 1). A block that covers the whole MSA was used for all positions in MSA-A, because it lacks gaps, whereas three blocks were generated for MSA-B, where the block width (*l*) at position 19 is one, (Fig. 1B, orange block). At position 19, the weights of the sequences not only of seq10a and seq10b but also of seq1-9 in MSA-B deviate drastically from those in MSA-A. Consequently, at position 19 of MSA-B, the weighted percentage of alanine, leucine, isoleucine and serine were equally 25 (Supplemental File 2). When *l* is one, the number of sequences which have *a* at *j* is *n_aj_* and the weighted probabilities of amino acids are 1/(*r_j_*∗*n_aj_*)\**n_aj_* = 1/*r_j_*. In MSA-A, those values were 62, 15, 12 and 11, respectively, demonstrating the limitation of PSI-BLAST PSSW calculation when the block width is tiny.

### Block extended PSI-BLAST (PSI-BLASTexB)

A simple and direct solution of this problem is to prevent block widths from being narrower than a certain width by exceptionally allowing gaps in the blocks. These gaps might cause the underestimation of gappy sequences in an alignment as discussed above, which however would certainly be a better estimation than the weights calculated for blocks having width of several residues.

The PSI-BLAST source code was downloaded from the BLAST FTP site (ftp://ftp.ncbi.nlm.nih.gov/blast/executables/blast+/2.5.0/ncbi-blast-2.5.0+-x64-linux.tar.gz). We revised the code of PSI-BLAST and added five lines after the line 1415 of ncbi-blast-2.5.0+-src/c++/src/algo/blast/core/blast_psi_priv.c as shown below.

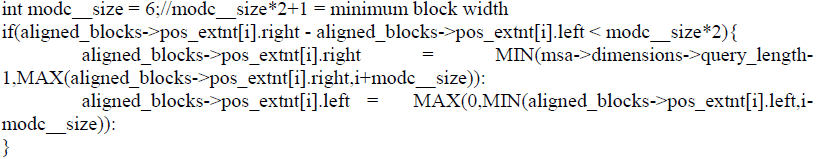

It implements the minimum block width (MBW), which is originally one. Blocks with widths < MBW are extended front and rear by MBW-1 until the termini of the MSA. For example, when MBW is 13, the deviated weights of MSA-B (Fig. 1B, red block) became similar to the weights of MSA-A (Fig. 1A). The resulting PSSM of MSA-B with MBW13 is provided as Supplemental File 3. The source code was configured with “––with-bin-release” and “––with-ncbi-public” options and compiled by the *make* command with no options. We designate the modified PSI-BLAST as PSI-BLASTexB.

### Benchmark dataset

The search performance was compared using SCOP20_training, SCOP20_validation and CATH20-SCOP datasets, which were established in our previous study (Yamada & Tomii 2014). The SCOP20_training and SCOP20_validation datasets were derived from the non-redundant set of 7074 proteins (SCOP20) which was provided by the ASTRAL compendium (Fox et al. 2014). The 7074 sequences were divided into two groups for parameter optimization (SCOP20_training) and performance evaluation (SCOP20_validation). CATH20-SCOP dataset was derived from the CATH database (Sillitoe et al. 2015) excluding sequences in the SCOP database. The sequences in the datasets were filtered so that the sequences did not have > 20% mutual sequence identity. Finally, our dataset included 3537, 3537, and 1754 sequences respectively. All datasets are available from http://csas.cbrc.jp/Ssearch/benchmark/.

### PSSM construction

PSSMs for individual sequences in the benchmark datasets were constructed using PSI-BLAST and PSI-BLASTexB against the Uniref50 dataset (Suzek et al. 2015) downloaded from the Uniprot FTP site (ftp://ftp.uniprot.org/pub/databases/uniprot/uniref/uniref50/). In this study, PSSMs for an iteration X were generated using the following command:

psiblast -query <QUERY> -db <UNIREF50 DB> -out_pssm <PSSM PATH> -num_iterations <X> -num_alignments 1000000.

We also extracted an MSA consisting of hits from a PSI-BLAST search with “-num_interation 1” option, and used the MSA directly to another search against the same dataset using the “-in_msa” option, which is an alternative means of running an iterative PSI-BLAST search using the MSA instead of the check point PSSM.

### Performance evaluation

Similarity searches were conducted against benchmark datasets using the constructed PSSMs and MSAs as queries using the “-in_pssm” and “-in_msa” options, respectively, instead of the “-query” option. We followed the rule set by Julian Gough (http://www.supfam.org/SUPERFAMILY/ruleset.html) (Gough et al. 2001) to judge true positive (TP) and false positive (FP) hits at the superfamily level. Superfamily definitions of the rule set differ from the original definitions of SCOP. The rule set also excludes hits with a potential homologous relation from FPs.

To evaluate the performance, we introduced an ROC plot, which has been widely used for performance evaluation (Angermuller et al. 2012; Biegert & Soding 2009). Hits from all queries were pooled and ranked by E-values. Then TP and FP hits until various E-value thresholds were counted and shown, with weighting of the TP and FP counts by 1/(number of all TPs in the dataset) for each query.

We also calculated the ROC5 score for hits with E-values less than 1.0, which indicates the search performance of individual queries using the following equation:

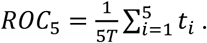

Therein, *T* signifies the total TP count; *t_i_* denotes the TP count until the *i*-th FP appears (Gribskov & Robinson 1996).

## Results and Discussion

We checked the distribution of block widths used for individual query positions at the second, third, fourth, and fifth iterations (Fig. 2). About 8%, 8%, and 5% of the blocks had widths of less than 5 amino acids (aa) at the fifth iteration of SCOP20_training, SCOP20_validation and CATH20-SCOP datasets, respectively. This fact indicates that the narrow-width blocks are constantly produced by PSI-BLAST.

**Figure 2.**
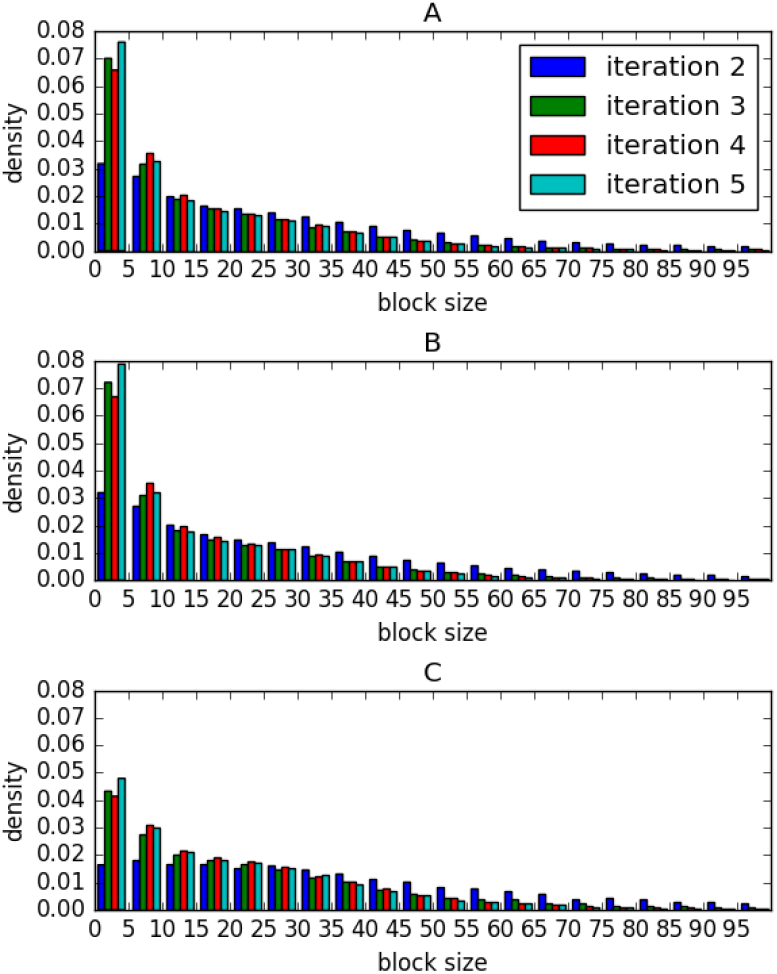
Distributions of block widths used for PSSW calculation with varying numbers of iterations. Results of searches against SCOP20_training, SCOP20_validation, and CATH20-SCOP are shown, respectively, in A, B, and C. Searches that converged before the fifth iteration were not used. Numbers of sequences (and blocks) used in A, B and C are, respectively, 2973 (532170), 2423 (438435), and 1351 (172450).

Using the SCOP20_training dataset, we analyzed the PSI-BLASTexB performance with varying MBW values (5, 13, 25 and 41) at the fifth iteration. PSI-BLAST corresponds to PSI-BLASTexB with the MBW of one. As Figure 3A shows, the performance of PSI-BLASTexB is much higher than that of PSI-BLAST across all MBW values. The performances are almost identical when MBW values are 13, 25, and 41, and are slightly low when MBW is 5, which suggests that 5 aa long blocks are insufficient to calculate the correct PSSW. Because the weighted TP count was highest when MBW was 25 at the false discovery rate (FDR) of 10%, we use the value as the default in the following experiments.

**Figure 3.**
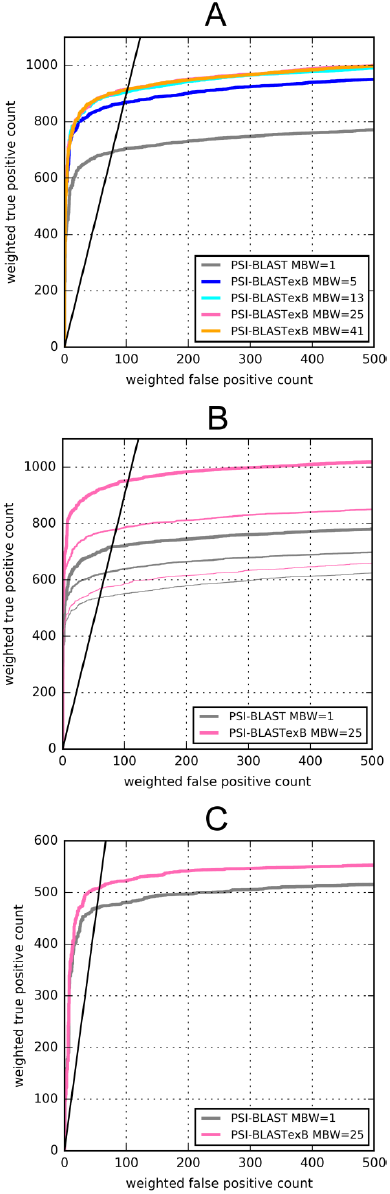
ROC curves of PSI-BLAST and PSI-BLASTexB. (A) ROC curves of PSI-BLAST (MBW=1) and PSI-BLASTexB (MBW = 5, 13, 25, or 41) at the fifth iteration against SCOP20_training. (B) ROC curves among searches with different numbers of iterations against SCOP20_validation. Narrow, normal, and thick lines respectively show the second, third, and fifth iterations. (C) ROC curves of PSI-BLAST and PSI-BLASTexB at the fifth iteration against CATH20-SCOP. Black lines represent FDR of 10%.

The performance improvement was also clear for SCOP20_validation and CATH20-SCOP (Figs. 3B and 3C). However, the performance improvement for CATH20-SCOP was slight compared with those of SCOP20_training and SCOP20_validation. That result is consistent with the result of the distributions of block widths. The fractions of narrow-width blocks in CATH20-SCOP are smaller than those of SCOP20_training and SCOP20_validation (Fig. 2), which is expected because our new method would be of no use if there were few narrow-width blocks.

To observe the relation between performance improvement and the block extension for each query, the incremental ROC5 scores (ROC5 score by PSI-BLASTexB - ROC5 score by PSI-BLAST) are shown against the ratio of positions with one aa long blocks at the second iteration for each query (Fig. 4). When the ratio is larger than 0.1, in other words, when more than 10% of PSSM positions are derived from one aa long blocks, 90, 85 and 81 PSI-BLASTexB searches among 182, 184 and 196 achieve higher performance than PSI-BLAST searches, and only for a 9, 11 and 9 cases PSI-BLASTexB searches are worse than PSI-BLAST searches against SCOP20_training, SCOP20_validation and CATH20-SCOP, respectively. In contrast, improvement of queries with a ratio less than 0.1 appears to be more random, though PSI-BLASTexB searches are also effective for many queries with a ratio less than 0.1. These results show how widening the widths of narrow blocks improves the search performance.

**Figure 4.**
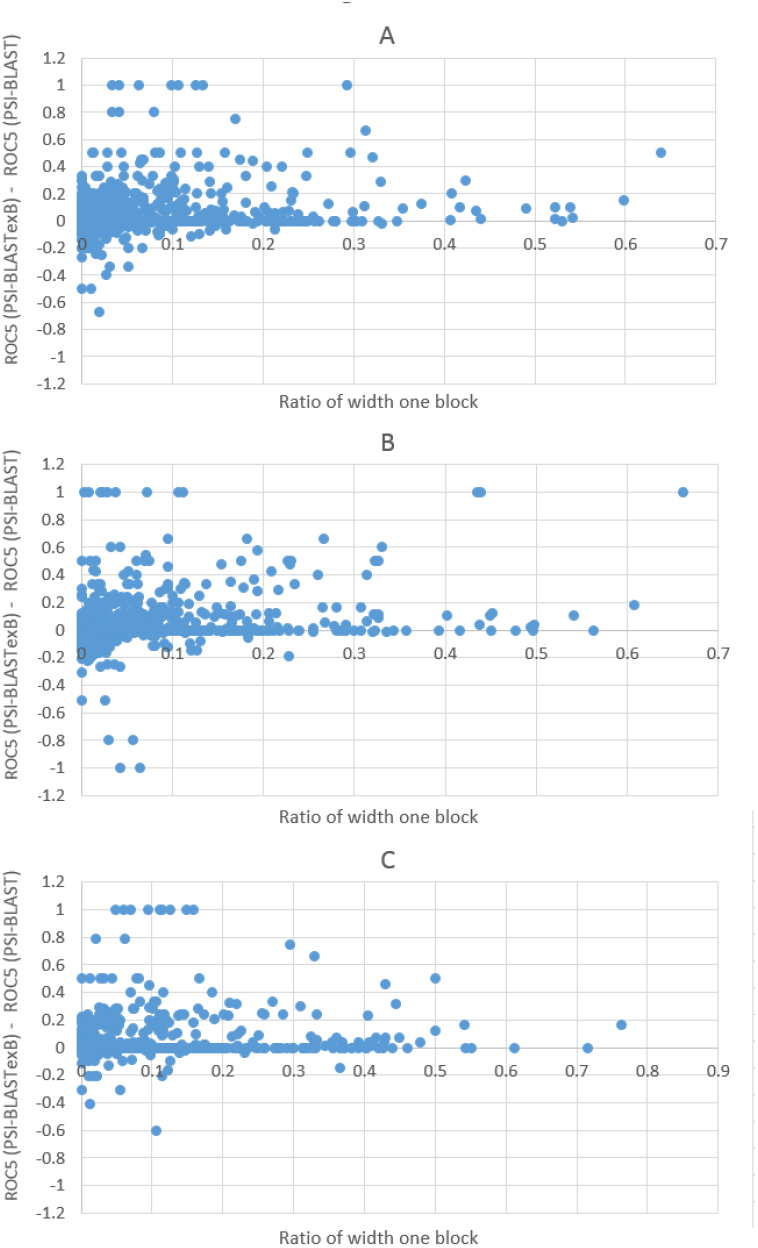
Relations between the ROC5 score improvement and the fraction of narrow blocks. The X-axis shows (number of one aa long blocks during PSSM construction)/(length of the query). The Y-axis shows the ROC5 score of PSI-BLASTexB replaced by that of PSI-BLAST. Each dot represents the result of a single query. Results of SCOP20_training, SCOP20_validation, and CATH20-ScOP at the second iteration are presented respectively in A, B, and C.

Recent versions of PSI-BLAST support a search using an MSA as an input with “-in_msa” option. We constructed MSAs from the outputs of PSI-BLAST and PSI-BLASTexB to use them as queries for the next search (see *Methods* for details). As Figure 5 shows, the performance of PSI-BLAST with “-in_msa” option is distinguishably lower than that of normal PSI-BLAST search with the corresponding number of iterations (Fig. 5). From our understanding, when “-in_msa” is used, PSI-BLAST divides a sequence in an MSA into multiple pieces if there are large gaps within (10 aa in case of ver. 2.5.0). Therefore, more narrow-width blocks are generated with the “-in_msa” option. Block extension by PSI-BLASTexB effectively suppresses performance degradation using MSAs as queries (Fig. 5). Therefore, PSI-BLASTexB can facilitate the use of MSAs prepared in advance as queries, e.g. Pfam seed alignments (Finn et al. 2016), HMM-HMM alignments by HHblits (Remmert et al. 2012), and progressive alignments by MAFFT (Katoh & Standley 2013) for distant homology detection.

**Figure 5.**
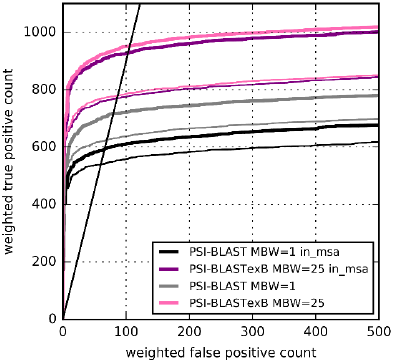
ROC curves with “-in_msa” option of PSI-BLAST and PSI-BLASTexB against SCOP20_validation. Thick and narrow lines respectively show ROC curves at the fifth and third iterations. The black straight line shows FDR of 10%.

Two factors can cause narrow-width blocks. One is a query dependent factor when there are multiple highly conserved regions in an MSA, such as multi-domain proteins, the blocks in boundaries (or close to boundaries) of those regions are more likely to become narrow. The other is library dependent. We showed in Figure 1B that fragmented sequences in libraries cause narrow blocks. Dividing queries such that each query has only one conserved region or removing fragmented sequences from library can be work-arounds to reduce the number of narrow-width blocks. However, the practical applications of these procedures might require further considerations (e.g., how to determine “conserved” and “fragmented“). Consequently, our adjusted method is a more realistic means of preventing the narrow-width blocks than controlling query-dependent and library-dependent factors.

## Conclusion

Because of sequence weighting scheme limitations, the PSI-BLAST performance has been penalized until now. We developed a customized PSI-BLAST, designated as PSI-BLASTexB, which solved such problems with extremely simple modification of the PSI-BLAST code. PSI-BLASTexB significantly outperformed PSI-BLAST. Therefore, it is expected to be useful not only for distant homology search, but also for many downstream methods that depend on PSI-BLAST with trivial effort. Binaries and source codes of PSI-BLASTexB (MBW = 25) are available at https://github.com/kyungtaekLIM/PSI-BLASTexB.

## Acknowledgments

We thank Dr. Yoshinori Fukasawa (AIST) for helpful discussions. This work was partially supported by the Platform Project for Supporting in Drug Discovery and Life Science Research (Platform for Drug Discovery, Informatics, and Structural Life Science) from the Japan Agency for Medical Research and Development (AMED).

